# Empirical noise-mean fitness landscapes and the evolution of gene expression

**DOI:** 10.1101/436170

**Authors:** Jörn M. Schmiedel, Lucas B. Carey, Ben Lehner

## Abstract

The effects of cell-to-cell variation (noise) in gene expression have proven difficult to quantify, in part due to the mechanistic coupling of noise to mean expression. To independently evaluate the effects of changes in expression mean and noise we determined the fitness landscapes in mean-noise expression space for 33 genes in yeast. The landscapes can be decomposed into two principal topologies: the fitness effects of protein shortage and surplus. For most genes, the fitness impact of sustained (mean) and short-lived (noise) deviations away from the expression optimum are linked and of similar magnitude. Sensitivity to both protein shortage and surplus creates a fitness landscape in which an ‘epistatic ratchet’ uncouples the evolution of noise from mean expression, promoting noise minimization. These results demonstrate that noise is detrimental for many genes and reveal non-trivial consequences of mean-noise-fitness topologies for the evolution of gene expression systems.

**Highlights:**

- Expression fitness landscapes in mean-noise space allow quantitative independent assessment of the effects of noise and mean expression on fitness for 33 genes
- Landscapes are described by a combination of just two principal topologies: fitness defects due protein shortage, or due to protein surplus
- Direct evidence that high expression noise is detrimental to fitness for 50% of genes
- Mean expression and noise have equivalent impact on organismal fitness
- Landscapes created by sensitivities to protein shortage and surplus facilitate independent evolution of gene expression noise via an epistatic ratchet

## Introduction

The mapping between genotype and phenotype determines how genetic variation affects phenotypes and how in turn genotypes evolve under natural selection. An important molecular phenotype for each gene is its expression level (Figure 1A). The production of proteins is tightly controlled at multiple regulatory levels. Protein levels do, however, show considerable variation, not only across genotypes and environments, but also among isogenic cells within the same environment and even in the same cell over time (Elowitz et al., 2002; Ozbudak et al., 2002). This non-genetic variation in protein production results from the stochasticity of chemical reactions as well as from the variable levels of regulators (Blake et al., 2003; Paulsson, 2004; Thattai and van Oudenaarden, 2001).

**Figure 1.**
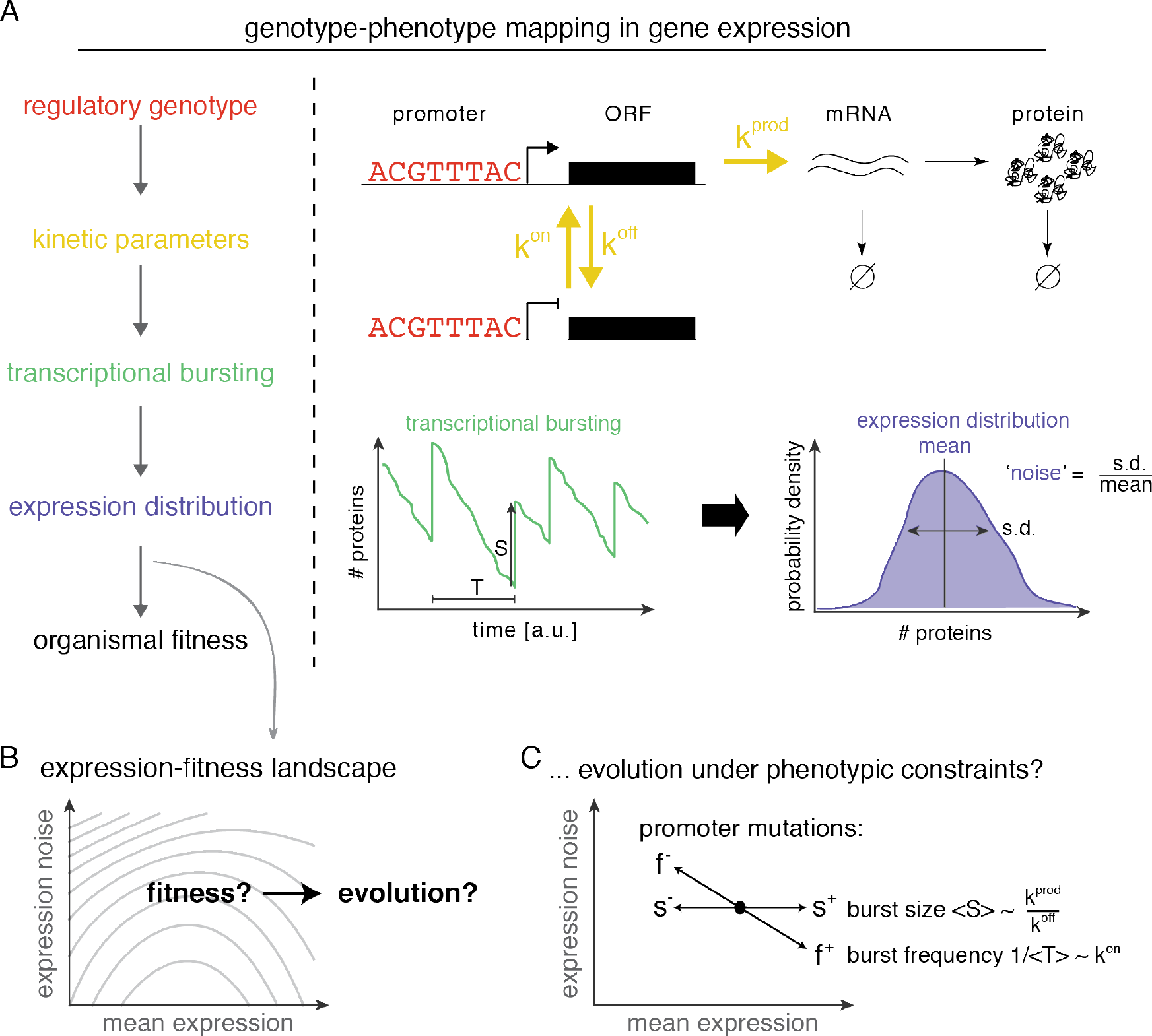
Genotype-phenotype mapping and the evolution of gene expression. (A) Multi-level mapping from genotype to expression phenotypes to organismal fitness. Regulatory genotypes (red) determine kinetic parameters (e.g. promoter on/off switching k^on^/k^off^, rate of mRNA transcription *k*^*prod*^, yellow), thus the properties of transcriptional bursting (average size <*S*> and frequency *1/<T>*, green), which results in particular expression distributions (blue). Expression distributions, characterized by their mean and noise (normalized standard deviation, ‘coefficient of variation’), determine organismal fitness via the expression-fitness landscape (panel B). (B) The expression-fitness landscape describes the mapping between the expression distribution of a gene (or its properties mean and noise) and fitness. Quantifying the topology of expression-fitness landscapes should inform about the evolution of gene expression systems. (C) The transcriptional process constraints how genetic variation can move gene expression in mean-noise space; mutations cannot lead to purely vertical moves. Does this affect the evolution of gene expression systems?

For any single environment and genotype, protein abundance in single cells therefore follows an expression distribution that is commonly characterized by its average (‘mean’) and its width (‘noise’). Understanding how these two properties impact fitness is important for understanding how regulation can evolve (Figure 1B).

However, the mean and noise of the expression distribution are not independent quantities, but are instead mechanistically coupled by the protein production process. In particular, switching between transcriptional permissive and prohibitive states leads to proteins being produced in bursts. While the size of bursts (the rates mRNAs and proteins are produced at when in the permissive state and how quickly genes revert back to a transcriptionally prohibitive state) only affects mean protein abundances, the frequency of bursts affects mean protein abundances and noise in an inversely proportional manner (Hornung et al., 2012; Pedraza and Paulsson, 2008; Raj et al., 2006; Raser and O’Shea, 2004; So et al., 2011). Mutations in promoters most often affect burst frequencies, resulting in anti-correlated changes in mean and noise (Hornung et al., 2012). Anti-correlations between mean abundances and noise are also observed comparing across genes (Bar-Even et al., 2006; Newman et al., 2006; Taniguchi et al., 2010).

Both large (Deutschbauer et al., 2005; Dietzl et al., 2007; Gerdes et al., 2003; Giaever et al., 2002; Hart et al., 2015; Hillenmeyer et al., 2008; Ramani et al., 2012) and small (Dekel and Alon, 2005; Dykhuizen et al., 1987; Keren et al., 2016; Rest et al., 2013) deviations of mean protein abundance from optimal levels can be detrimental to organismal fitness.

The fitness effects of noise are, however, less well explored. The consequences of changes in noise will depend on two factors (Duveau et al., 2018; Tanase-Nicola and ten Wolde, 2008): whether deviations from optimal protein abundances in a gene are detrimental to fitness and how close the mean protein abundance is to the optimal level.

If mean protein abundance is far from the optimal level high noise can be beneficial, allowing some cells to transiently express more optimal protein abundances (Duveau et al., 2018). In fluctuating environments, high expression noise may therefore be a ‘bet-hedging’ strategy to diversify phenotypes (Acar et al., 2008; Blake et al., 2006; Eldar et al., 2009; Maamar et al., 2007; Süel et al., 2007).

If mean protein abundance is, however, close to its optimum, as is presumably the case for many genes in a constant environment (Keren et al., 2016), then high noise is detrimental because fluctuations result in sub-optimal protein abundance. The observed low noise levels of many dosage-sensitive genes in yeast provides circumstantial evidence that noise is detrimental and has been selected against (Bar-Even et al., 2006; Batada and Hurst, 2007; Fraser et al., 2004; Hornung et al., 2012; Keren et al., 2016; Lehner, 2008; Newman et al., 2006; Wang and Zhang, 2011). However, the mechanistic coupling of noise and mean levels in protein production has made it difficult to directly test the fitness consequences of changes in expression noise alone. Notably, the Wittkopp lab has recently demonstrated that yeast strains in which the TDH3 gene, for which non-optimal abundances are detrimental to yeast fitness (Duveau et al., 2017), is driven by high noise promoters are less fit (Duveau et al., 2018). Consequently, noise-increasing mutations in its endogenous promoter have been found to be under purifying selection (Metzger et al., 2015).

Whether these results generalize to other genes is, however, unclear. Importantly, we still lack quantitative experimental data on the fitness effects of changes in noise and their interaction with the effects of changes in mean (Figure 1B). Therefore, how these two expression phenotypes might co-evolve, especially given their mechanistic couplings by the transcriptional process, is still an open question (Figure 1C).

Here we reconstruct fitness landscapes in mean-noise expression space for 33 genes in yeast using published fitness data of yeast strains in which genes are driven by a library of synthetic promoters (Keren et al., 2016; Sharon et al., 2014) (Figure 2A-C). These continuous landscapes allow for a comprehensive, quantitative assessment of both the independent as well as interdependent fitness effects of noise and mean expression. Principal component analysis of mean-noise-fitness landscapes reveals that all landscapes can be mapped back to just two principal landscape topologies, representing a gene’s sensitivity to protein shortage, and to surplus. These two principal topologies link the fitness effects of mean deviations and noise and thus determine how intolerant a gene is to high expression noise. Overall, half of the assayed genes are noise intolerant with evidence that high noise has been selected against during evolution. We further use the expression-fitness landscapes to explore how mean and noise can evolve, given their mechanistic coupling imposed by the transcriptional process. We find that on landscapes of genes sensitive to both protein shortage and surplus a specific topology, the noise funnel, uncouples the evolution of noise from mean via an epistatic ratchet-like mechanism, and therefore allows for the independent minimization of noise levels.

**Figure 2.**
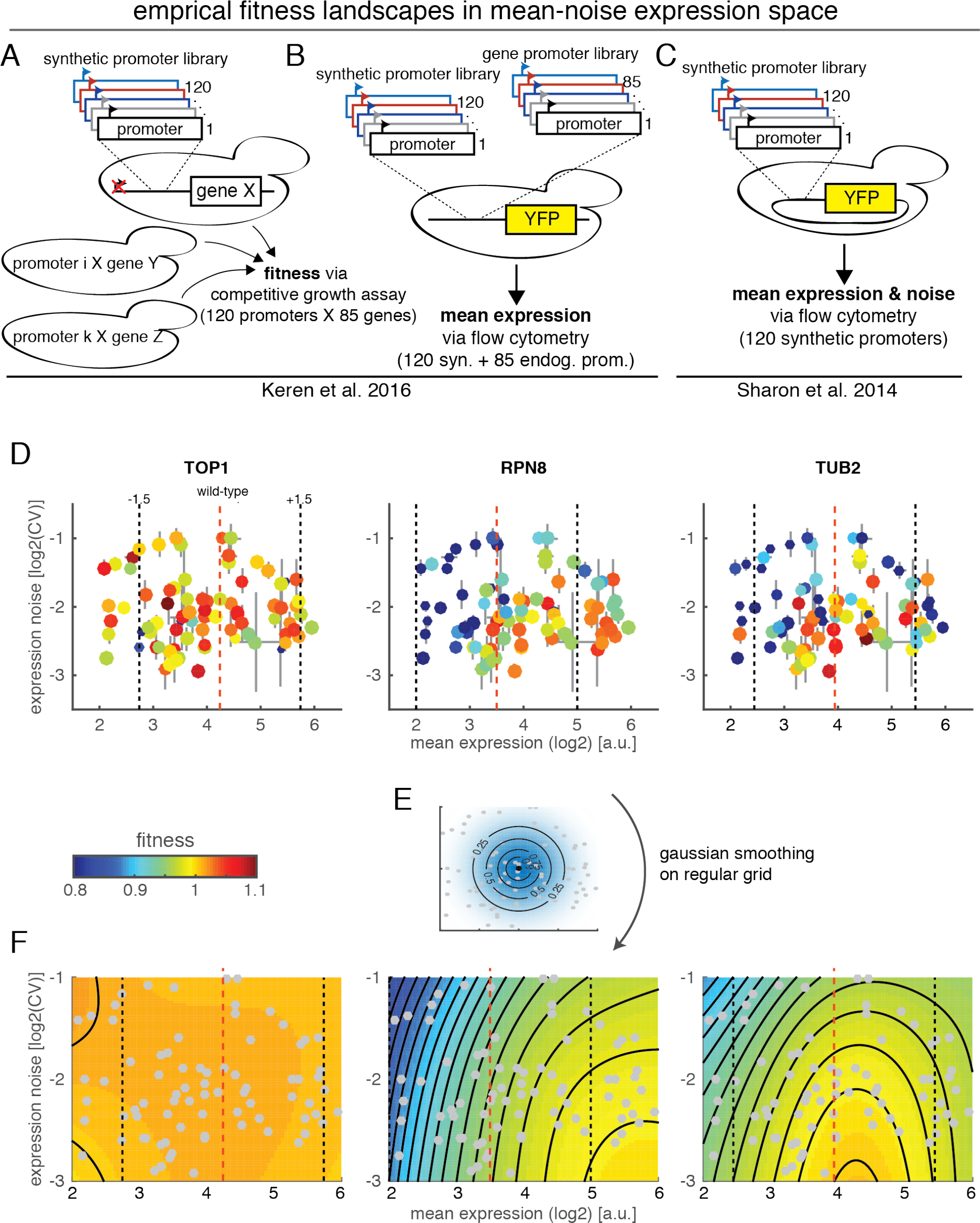
Noise-mean-fitness landscapes reveal gene-specific responses to expression deviations. (A) Keren et al. (2016) constructed yeast strains where in each strain one of a panel of genes (X, Y, Z,…) is driven by one of a panel of 120 synthetic promoters. All constructed strains were pooled and their fitness was measured in competitive growth experiments. (B) Relative mean expression strength of the synthetic promoters and the endogenous promoters of all investigated genes was measured by cloning them in front of YFP in the HIS3 locus and assaying the resulting strains individually by flow cytometry (Keren et al., 2016). (C) Sharon et al. (2014) measured mean expression and noise of the synthetic promoters via flow cytometry. Synthetic promoters were cloned in front of YFP on a plasmid. Fluorescent activated cell sorting in bins across the expression range in combination with deep sequencing was used to reconstruct expression distributions (and from these mean and variance/noise) of individual promoters. (D) Fitness of synthetic promoter-gene strains plotted in mean expression - expression noise (“mean - noise”) space for three example genes. Each panel shows the fitness of 79 yeast strains, in which a particular synthetic promoter drives the indicated gene, as a function of mean expression and expression noise of the synthetic promoters. Colour of circles indicates fitness, size of circles indicates measurement error of fitness measurement, error bars in horizontal and vertical directions indicate measurement error in mean and noise direction, respectively. Dashed vertical line shows estimated wild-type expression of the gene, red vertical lines mark region ±1.5-fold from wild-type expression (the region used in subsequent analyses). (E) Construction of noise-mean-fitness landscapes using Gaussian smoothing on a regular grid. Fitness of each grid point is calculated as weighted sum over the fitness of all strains, with weights depending on distance to grid point (shown) and measurement errors in all three dimensions (see Methods). (F) Smoothed noise-mean-fitness landscapes of the three genes shown in panel (D). Surface colour indicates fitness. Contour lines are spaced by 1% fitness. Strains (grey points), wild-type expression (red vertical line) and +− 1.5-fold around wild-type expression (grey vertical lines) are indicated. CV: coefficient of variation (standard deviation/mean of expression distribution).

Together, our analyses reveal the quantitative fitness effects of expression noise and their relation to mean expression and how the evolution of gene expression is shaped by phenotypic restrictions and expression-fitness landscape topology.

## Results

### Reconstruction of empirical fitness landscapes in expression mean-noise space

We obtained data on the fitness of yeast strains where in each strain one of a panel of genes is driven by one of a panel of 120 synthetic promoters (Keren et al., 2016). Here, in one set of experiments, the library of 120 synthetic promoters was cloned upstream of each of 85 open reading frames, replacing the endogenous promoter (Figure 2A). All constructed strains were pooled and their fitness was measured in competitive growth experiments. In a second set of experiments, the synthetic promoters as well as the endogenous promoters of all investigated genes were cloned in front of YFP in the HIS3 locus and flow cytometry was used to determine their relative average expression strength (Figure 2B). Together, this allowed the authors to analyze the fitness effects of mean expression changes relative to the wild-type expression of genes. In addition to this dataset, we also obtained data from an earlier study from the same group of authors that measured both mean and cell-to-cell variation (noise, coefficient of variation, standard deviation/mean) in expression of the same synthetic promoters driving YFP on a plasmid (Figure 2C) (Sharon et al., 2014).

While the absolute expression strength and noise of a particular promoter can depend on its genomic location, for the following analyses we make the assumption that the *relative* expression strength and noise levels between promoters is independent of the genomic location. The validity of this assumption is supported by the literature (Chen and Zhang, 2016; Schikora-Tamarit et al., 2018) as well as by the high correlation of mean expression strengths when the synthetic promoters are driving YFP from a plasmid or the HIS3 locus (R^2^ = 0.93, R^2^ = 0.99 after exclusion of 11 outliers, see Figure 2 - figure supplement 1A and Methods).

We filtered the set of promoters used in the original studies according to several quality control criteria and in order to obtain a homogenously populated region in the expression mean - noise space (Figure 2 - figure supplement 1, Methods). In the final dataset each gene is expressed under the control of 74 to 79 (average 78) different synthetic promoters that span an expression range of ~16-fold and a noise range of ~4-fold, as assessed by the coefficient of variation (Figure 2D).

To systematically study the fitness effects of varying both mean and noise around wild-type expression levels we restricted our analyses to the 33 genes with wild-type expression levels in the centre of the well-populated mean-noise space (Figure 2 - figure supplement 1D, see Discussion for consideration of effects away from wild-type levels). These genes represent a wide range of cellular functions, such as transcription factors, RNA polymerase, proteasome, cytoskeleton, trafficking and metabolism.

We started our analysis by examining the fitness of gene-promoter strains in the expression mean-noise space (Figure 2D). For some genes, like the topoisomerase I TOP1, all strains across the mean-noise space have approximately wild-type fitness. In contrast, for the 26S proteasome subunit RPN8 strains with low expression/high noise promoters tend to have low fitness. In addition to low fitness at low expression/high noise, for the beta-tubulin TUB2 also strains with high expression/high noise promoters display lowered fitness.

We sought a systematic way to investigate how mean and noise impact fitness, both together and independently. We reasoned that for each gene there exists a continuous fitness landscape in the mean-noise expression space. This landscape has been experimentally sampled by the different synthetic promoter strains.

To reconstruct a smooth, continuous fitness landscape for each gene we calculated fitness values on a regular grid across the mean-noise space using a Gaussian smoothing approach. For each point on the gird a fitness value was calculated as the weighted sum of all measured fitness values for that gene. Weights were calculated according to a bivariate normal kernel (Figure 2E), centred on the grid-point and with gene-independent scaling parameters in mean and noise direction optimized to minimize the root mean squared error between the smoothed fitness landscapes and the raw data (estimated using ten-fold cross-validation). The weighting of each synthetic promoter strain was further modified by the measurement error of its mean expression, noise and fitness values (see Methods for full details).

The reconstructed mean-noise fitness landscapes reveal that for TOP1 there is essentially no systematic effect of mean expression or expression noise on fitness (Figure 2F, see Figure 3 for all landscapes). The fitness landscape of RPN8 reveals coupled negative fitness effects of lowered mean expression and high noise. Finally, the fitness landscape of TUB2 reveals a non-linear relationship between mean and noise on fitness. High expression noise is always detrimental, but the effect of noise on fitness increases as mean expression deviates more from the wild-type expression level.

**Figure 3.**
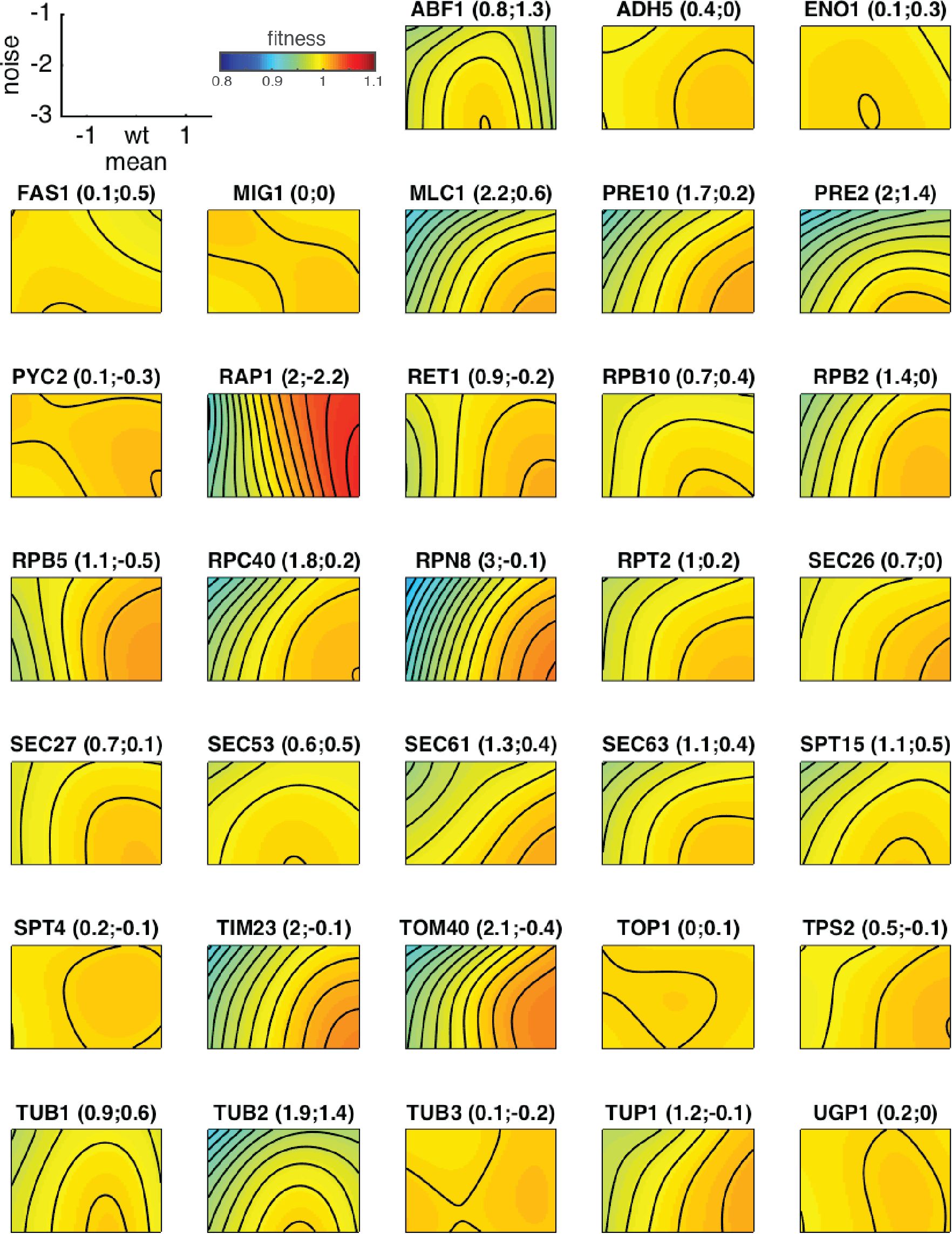
Expression-fitness landscapes in mean-noise space around wild-type mean expression. All 33 expression-fitness landscapes in mean-noise space ±1.5 log2-units around their wild-type mean expression of all 33 genes. Colours indicate fitness levels; contours are spaced in increments of 0.01 fitness units. Plot titles give gene name and principal topology 1 and 2 loadings, respectively (related to Figure 4A).

Together, expression-fitness landscapes in the mean-noise space present a valuable opportunity to study the interplay of two molecular phenotypes of gene expression on fitness, in a quantitative and systematic manner.

### The principal topologies of expression fitness functions

To systematically study the noise-mean fitness landscapes, we first investigated whether there are any commonalities between the fitness landscapes of all genes by performing a principal component analysis across all landscapes using the 8-fold mean expression range around the predicted wild-type expression of each gene (Figure 4 - figure supplement 1A).

Strikingly, the principal component analysis revealed two dominant topologies, together which explain 96% of the variance across landscapes (Figure 4A and Figure 4 - figure supplement 1B).

Common to both principal topologies are their intolerance for high expression noise (Figure 4F). Moreover, both topologies show a monotonically saturating relationship between fitness and protein abundance, they do however differ in the directionality of this relationship (Figure 4G).

**Figure 4.**
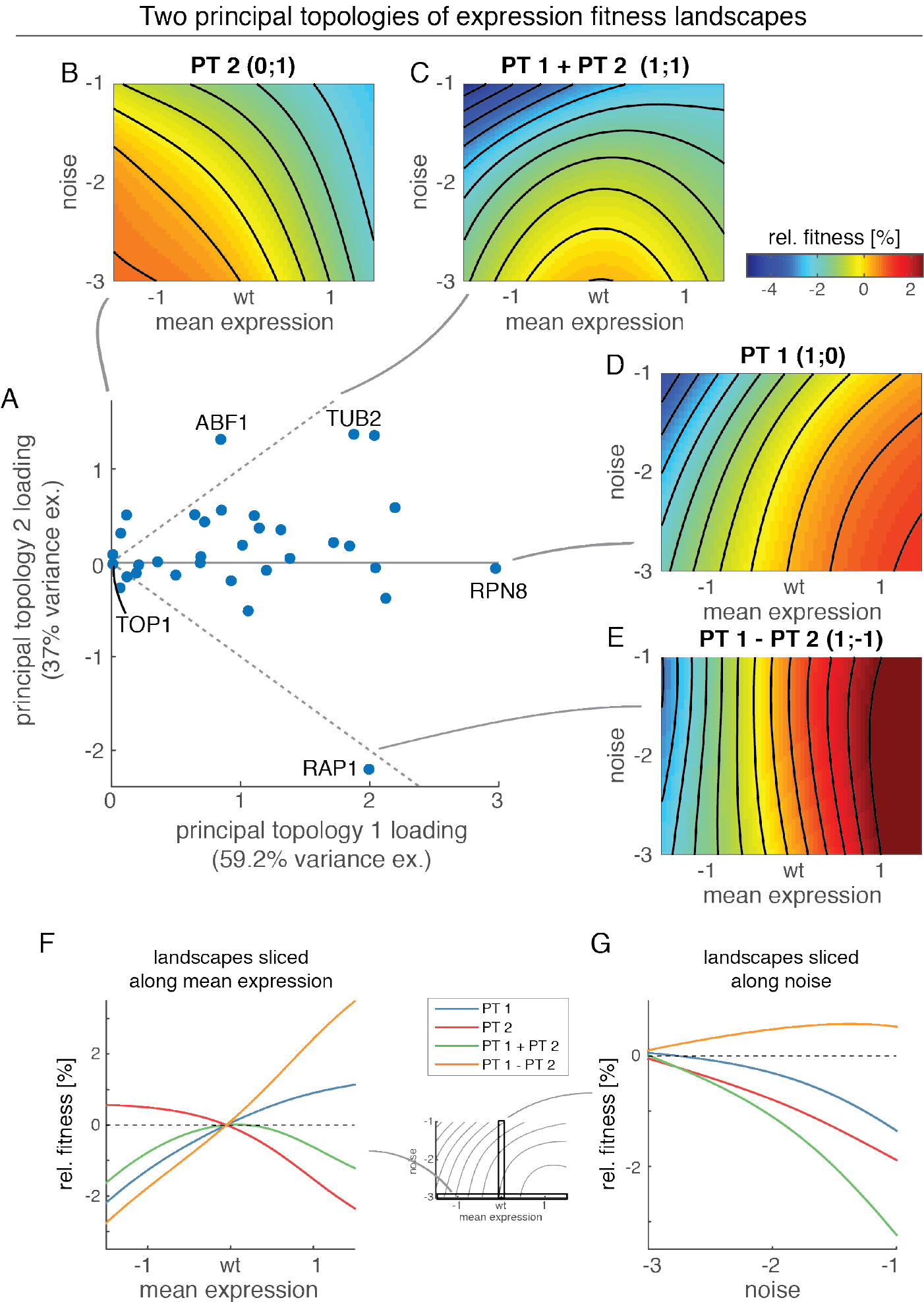
Two principal topologies of expression fitness landscapes. (A) Principal component analysis of expression fitness landscapes reveals two principal topologies (PT) that explain the majority of variation observed across all landscapes. Scatter plot shows PT loadings of individual landscapes (dashed lines show equivalent loadings with both PTs). (B-E) Landscapes show individual principal landscape topologies (PT1 and PT2) and their equivalent combinations (PT1 + PT2 and PT1 - PT2). (F) Principal topologies sliced along mean expression range at minimal expression noise (log_2_(CV) = −3). (G) Principal topologies sliced along noise range at wild-type mean expression.

The first principal topology exhibits high fitness if mean expression is at or above wild-type mean expression and if expression noise is low (Figure 4D). Fitness drops, however, for both lower than wild-type mean expression and higher noise. The first principal topology therefore represents the fitness consequences of protein shortage.

In contrast, the second principal topology has high fitness at or below wild-type mean expression and at low expression noise, but lower fitness at high mean expression or high noise (Figure 4B); it therefore represents the fitness consequences of protein surplus.

Individual landscapes are made up of different combinations of the two principal topologies (Figure 4A). All landscapes have a non-negative loading for the first principal topology, suggesting that the fitness effects of protein shortage are at best neutral but are detrimental for most genes. Indeed, loadings for the first principal topology are predictive of a gene’s essentiality (Figure 4 - figure supplement 1C).

In contrast, genes show both positive as well as negative loadings for the second principal topology. Combinations of positive loadings for both topologies lead to peaked landscapes, with decreased fitness and amplified negative impact of high noise when mean expression deviates from wild-type expression in either direction (Figure 4C). Three genes (ABF1, TUB2 and PRE2) show pronounced peaked patterns, consistent with findings of essentiality as well as sensitivity to copy number amplifications for all three genes (Giaever et al., 2002; Makanae et al., 2013; Sopko et al., 2006). An additional nine genes (MLC1, RPB10, RPT2, SEC27, SEC53, SEC61, SEC63, SPT15, TUB1) show somewhat weaker peaked patterns. While eight of these genes are essential for growth, none of these genes has previously been found to be sensitive to overexpression, suggesting that the patterns observed here might be subtler than those that can be detected by large-scale overexpression screens (Makanae et al., 2013; Sopko et al., 2006).

On the opposite end of the scale, one gene, RAP1, shows a significant negative loading for topology 2 that is as large as its positive loading for topology 1. The result is a landscape that shows a continuous increase in fitness with higher protein expression and no intolerance of expression noise at all (Figure 4E). RAP1 is known to have varied transcriptional effects on many yeast genes (Lieb et al., 2001), including the activation of both ribosomal proteins and glycolytic enzymes (Brindle et al., 1990; Shore and Nasmyth, 1987), and we hypothesize that its mild overexpression in the specific glucose-rich growth conditions might thus lead to increases in growth rate, even though strong overexpression of RAP1 is toxic (Freeman et al., 1995; Sopko et al., 2006). It is unclear, however, why RAP1 might be completely insensitive to varying expression noise levels.

In summary, two principal topologies in mean-noise expression space - representing the elemental response to having too few or too many proteins - explain nearly all variability in the reconstructed fitness landscapes. This suggests a hard-wired quantitative relationship between how fitness is affected by short-term (noise) and sustained (mean) deviations from optimal protein abundance.

### Noise intolerance is related to expression sensitivity and predicts endogenous protein expression noise

We next statistically quantified the effects of mean and noise on fitness and their relationship across fitness landscapes. We calculated for each gene the effect of mean expression changes on fitness, its *expression sensitivity*, as the average curvature of the mean expression-fitness function close to wild-type mean expression at minimal expression noise levels (log_2_(CV) = −3) (Figure 5A) (Keren et al., 2016). Additionally, we quantified for each gene the fitness effect of expression noise, its *noise intolerance*, as the maximal negative slope of the noise-fitness function at wild-type mean expression (Figure 5B). Notably, assessment of *noise intolerance* is robust to the exact metric chosen (Figure 5 - figure supplement 1A).

**Figure 5—.**
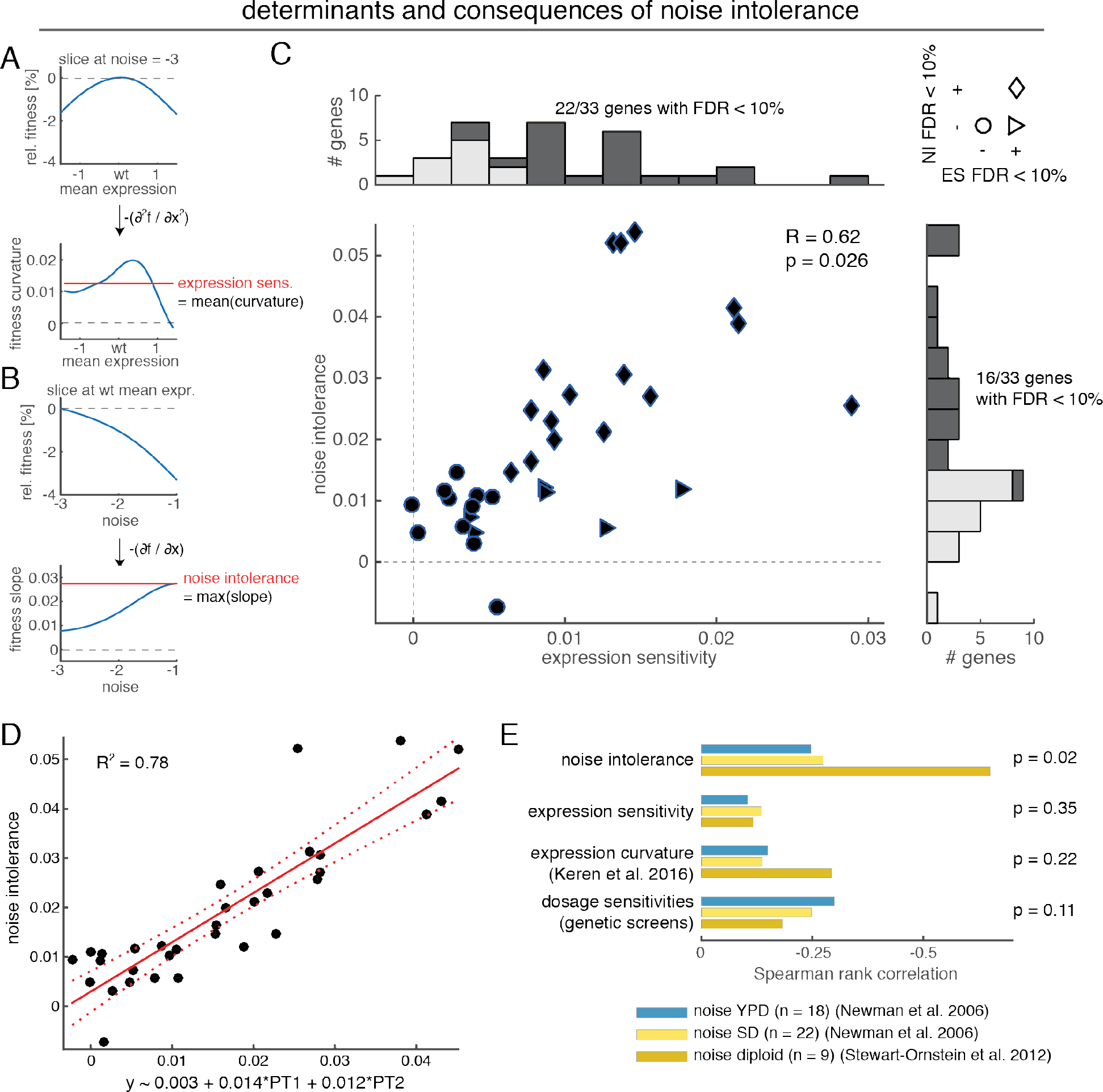
Noise intolerance correlates with expression-sensitivity and predicts endogenous noise levels. (A) Expression sensitivity is calculated as the average curvature of the fitness function ±1-fold around wild-type expression. The curvature of the fitness function is the second partial derivate of fitness with respect to mean expression at low noise (log_2_(CV) = −3). (B) Noise intolerance is calculated as the maximum slope of fitness with respect to increased noise. The slope is calculated as the negative partial derivative of fitness with respect to noise at wild-type mean expression. (C) Relationship between expression sensitivity and noise intolerance across all genes. False discovery rate (FDR) < 10% threshold indicated in both dimensions. Histograms on top and to the right show distributions of genes across expression sensitivity and noise intolerance, respectively. Indicated are the numbers of genes with FDR < 10%. Pearson correlation coefficient and p, the fraction of 10^4 sets of randomized landscapes that have Pearson correlation coefficients greater than the one found on the real landscapes, are indicated. (D) Linear model to predict noise intolerance from principal topology loadings (PT1: principal topology 1 loading, PT2: principal topology 2 loading). Red solid line shows best fit, dashed lines show 95% confidence interval. X-axis label gives linear model coefficients. (E) Spearman rank correlation between predictors for genes’ intolerance of noise (noise intolerance and expression sensitivity from this study, curvature of expression-fitness curve (Keren et al., 2016), dosage sensitivity of gene estimated from large-scale screens) and three sets of endogenous noise measurements (n: number of genes with available noise measurements). P-value given is the aggregated p-value from Fisher’s method across all three datasets.

Two thirds of the assayed genes (22 out of 33) are significantly expression sensitive (at false discovery rate (FDR) < 10%, estimated using randomized controls), i.e. for these genes deviations from wild-type mean expression leads to measurably reduced fitness (Figure 5C). Consistently, expression sensitivity estimated from fitness landscapes is highly predictive of known dosage sensitivities assessed from large-scale deletion or overexpression screens (Figure 5 - figure supplement 1B).

Moreover, half of all genes (16 out of 33) are significantly noise intolerant (at FDR < 10%, estimated using randomized controls), i.e. for these genes higher noise at wild-type expression levels leads to reduced fitness. These results are therefore rare direct evidence based on experimental data that high expression noise, i.e. short-lived expression fluctuations away from optimal wild-type expression, does impact organismal fitness for many genes in yeast.

In line with our findings from the principal topologies of fitness landscapes and previous reasoning (Bar-Even et al., 2006; Batada and Hurst, 2007; Fraser et al., 2004; Lehner, 2008; Newman et al., 2006), expression sensitivity and noise intolerance are correlated across genes (Pearson correlation coefficient R = 0.62). Importantly, this correlation does not arise from how we have reconstructed fitness landscapes (p = 0.026 in comparison to 10^4^ sets of random landscapes, Figure 5C). Genes that are sensitive to changes in mean expression levels thus also tend to be more intolerant to high gene expression noise.

Similar conclusions, in terms of the relationship between expression sensitivity and noise intolerance as well as the significance of their effects, are reached if both measures are instead estimated from partial correlations on the raw data of gene-promoter strains (Figure 5 - figure supplement 1C).

Moreover, we find that in a linear model to predict noise intolerance from the two principal topologies, both topologies contribute with nearly equal weights to explain a maximum of 78% of the variance in noise intolerance (Figure 5D, weights and standard error: 0.014 ± 0.002 and 0.012 ± 0.002 for first and second topology, respectively); suggesting that sensitivity to both too few as well as too many proteins contributes to intolerance of fluctuations in protein abundances.

Together with previous analyses (Bar-Even et al., 2006; Batada and Hurst, 2007; Duveau et al., 2018; Fraser et al., 2004; Lehner, 2008; Newman et al., 2006), these results suggest that too much noise impairs fitness for many yeast genes. During evolution, therefore, selection may have acted to minimize noise in the expression of these noise intolerant genes. To test this, we compared how the noise intolerance quantified on each genes’ fitness landscape relates to its measured *in vivo* protein expression noise in multiple published datasets (Figure 5E). Noise intolerance is indeed negatively correlated with the endogenous protein noise level of genes in three different large-scale datasets (Spearman rank correlation: ρ = −0.25 (noise in YPD), ρ = −0.27 (noise in SD), ρ = −0.65 (noise diploids), aggregated p-value from Fisher’s method, p = 0.02) (Newman et al., 2006; Stewart-Ornstein et al., 2012). Moreover, noise intolerance is a better predictor of endogenous protein noise levels than expression-sensitivity estimated from the fitness landscapes (ρ = −0.1, ρ = −0.14, ρ = −0.12; p = 0.35), the expression curvature of expression-fitness functions as previously estimated (ρ = −0.15, ρ = −0.14, ρ = −0.29; p = 0.22)(Keren et al., 2016) or known dosage sensitivity of genes (ρ = −0.3, ρ = −0.25, ρ = − 0.18; p = 0.11).

Together, this provides good evidence that selection has acted during the evolution of budding yeast to minimize fluctuations in gene expression due to their detrimental impact on organismal fitness.

### Fitness is similarly sensitive to changes in mean expression and noise

We next sought a more detailed view of how fitness reacts to changes in noise and mean expression across the landscapes, in order to explore how gene expression systems might evolve. Across three stereotypical landscapes (the two principal topologies and their combination, the ‘peaked’ landscape) we calculated the fitness slopes in noise and mean directions on various points on the landscapes. The resulting vectors indicate the sensitivity of fitness to changes in noise or mean expression and generally point in the direction of the fitness peak on the landscapes (Figure 6A).

We first analysed the impact of noise and mean changes on fitness dependent on the absolute fitness at a location on the landscapes. We find that the magnitude of slopes depends on the absolute fitness at locations on the landscape (Figure 6B). That is, changes in mean and noise have smaller impacts on fitness if fitness is already high, which is consistent with the saturating relationships between fitness and both mean expression and noise observed (see Figures 4F and G).

**Figure 6.**
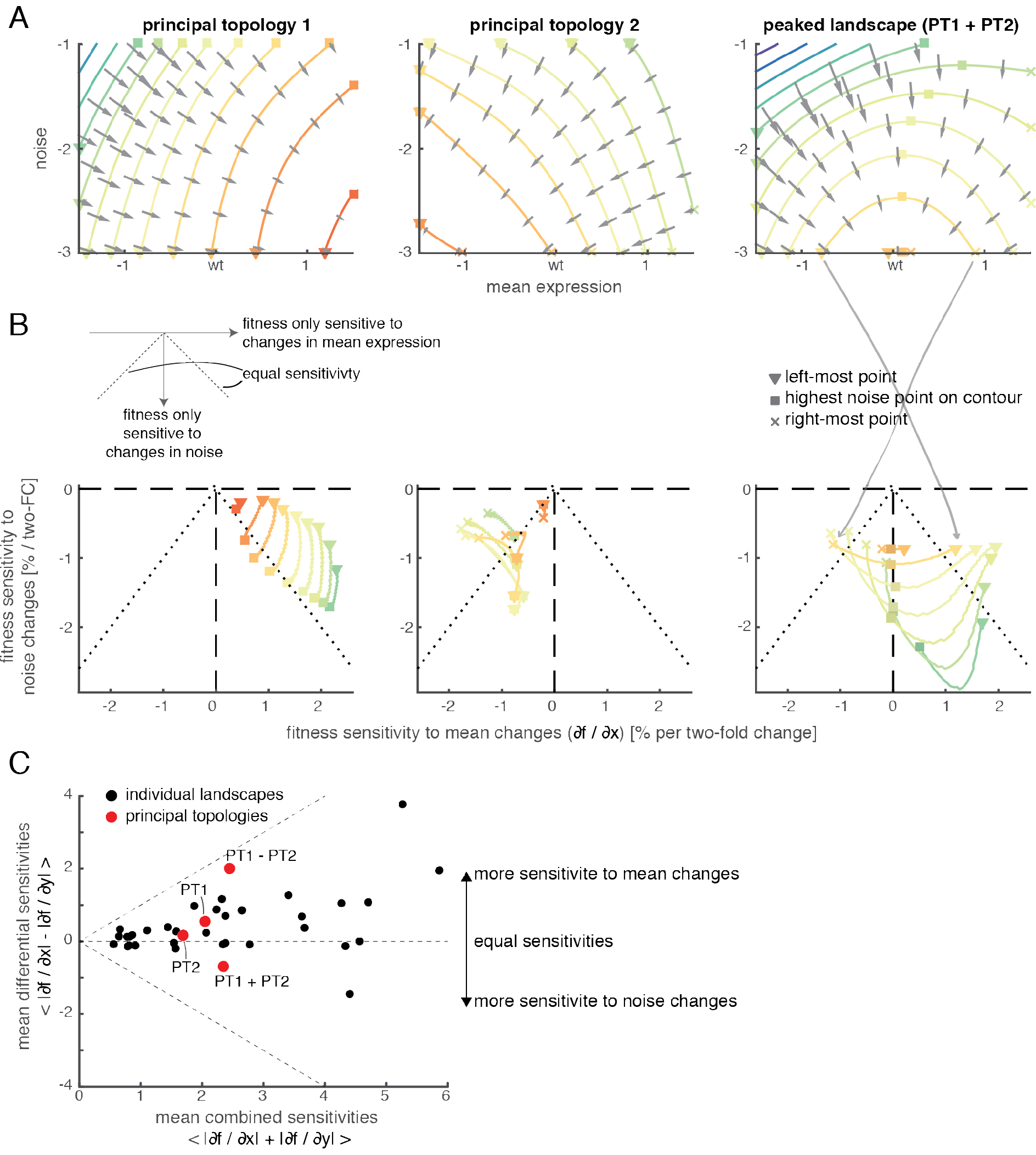
Analysis of fitness sensitivities to changes in mean expression and noise reveals similar magnitudes and evolutionary contingency. (A) Fitness sensitivity towards changes in mean and noise on principal topologies (left and middle) and peaked (topologies 1+2, right) landscapes. Sensitivities are calculated as slopes of the fitness surface in mean or noise direction (normalized to give the percent fitness change upon a two-fold change in mean or noise). Vectors show the combination of slopes in both directions. Contours show levels of equal fitness (“equi-fitness lines”, fitness colour scale as in Figure 4A). Triangle indicates left-most, square indicates ‘highest-noise’ and ‘x’ indicates right-most (if applicable) point on contour lines. (B) Fitness sensitivities to changes in mean expression and expression noise directions on contour lines from panel (A) (fitness slopes in mean or noise directions). Colours according to fitness of contour lines as in panel (A). Shapes indicate locations on contour lines as indicated in panel (A). (C) Comparison between fitness sensitivities to changes in mean expression and expression noise across principal topologies and individual gene landscapes. Shown are combined (summed) or differential absolute fitness sensitivities in both directions averaged across all grid-points on each landscape. Principal topologies and their combinations are highlighted in red. Diagonal dashed lines show where landscapes are only sensitive to changes in mean (upper) or noise (lower), middle line indicates equal sensitivities to changes in mean and noise.

Moreover, the absolute impacts of changes in mean or noise on fitness at any point on the stereotypical landscapes landscape are correlated and of similar magnitude (Figure 6B), suggesting that reducing expression noise is as important as optimising mean expression in order to maximize organismal fitness. To test whether this also applies to individual gene landscapes, we compared fitness sensitivities to changes in mean and noise on each grid-point across all individual landscapes. Indeed, on average, the impact of changes in mean or noise on fitness are similar across all landscapes (Figure 6C).

### Differential fitness sensitivity to changes in mean expression and noise creates contingency on fitness landscapes

Next, we examined how the sensitivities to changes in mean and noise are related across points on the landscape that have similar fitness, in order to understand whether evolution on fitness landscapes is contingent. When comparing sensitivity along such equi-fitness contour lines, we find that sensitivities to changes in mean or noise tend to be anti-correlated (Figure 6B).

On one hand, fitness is more sensitive to changes in mean expression when expression noise is low (and fitness therefore insensitive to changes in noise). Therefore, changes in mean expression are more likely to be selected for or against if a gene has low expression noise.

On the other hand, fitness is most sensitive to changes in noise on the highest noise locations of equi-fitness lines. This is most apparent on peaked expression-fitness landscapes (PT1+PT2), where these noise maxima of equi-fitness lines coincide with mean expression being optimal and therefore fitness is insensitive to changes in mean expression. However, because phenotypic change in noise is coupled to changes in mean expression via the transcriptional process, this poses the question of how noise could possibly evolve towards lower levels from these locations in the landscape when mean expression is already optimal.

### A noise funnel on peaked landscapes uncouples the evolution of noise from mean expression

We therefore next investigated how gene expression would evolve under the phenotypic restrictions on changes in noise and mean expression imposed by the transcriptional process.

In gene expression, mutations in regulatory elements, e.g. the promoter region, have specific effects on mean expression and expression noise that are determined by how they affect the underlying molecular mechanisms of transcriptional bursting (Hornung et al., 2012; Pedraza and Paulsson, 2008; Raj et al., 2006; Raser and O’Shea, 2004; So et al., 2011) (Figure 7A). Mutations can affect the *frequency* of transcriptional bursts, for example by affecting the rate with which transcription factors bind the promoter. An increased frequency of transcriptional bursts leads to higher average number of mRNAs produced but also to less noise in the levels of mRNAs. Or mutations can affect the *size* of transcriptional bursts, for example by affecting the rate at which transcription factors dissociate from the promoter or by altering the rate at which mRNAs are produced when the promoter is active. Changes in burst size alter mean expression but do not change expression noise. The molecular mechanisms underlying the transcriptional process and how genetic variability affects them thus restrict how genes can evolve on expression fitness landscapes.

**Figure 7.**
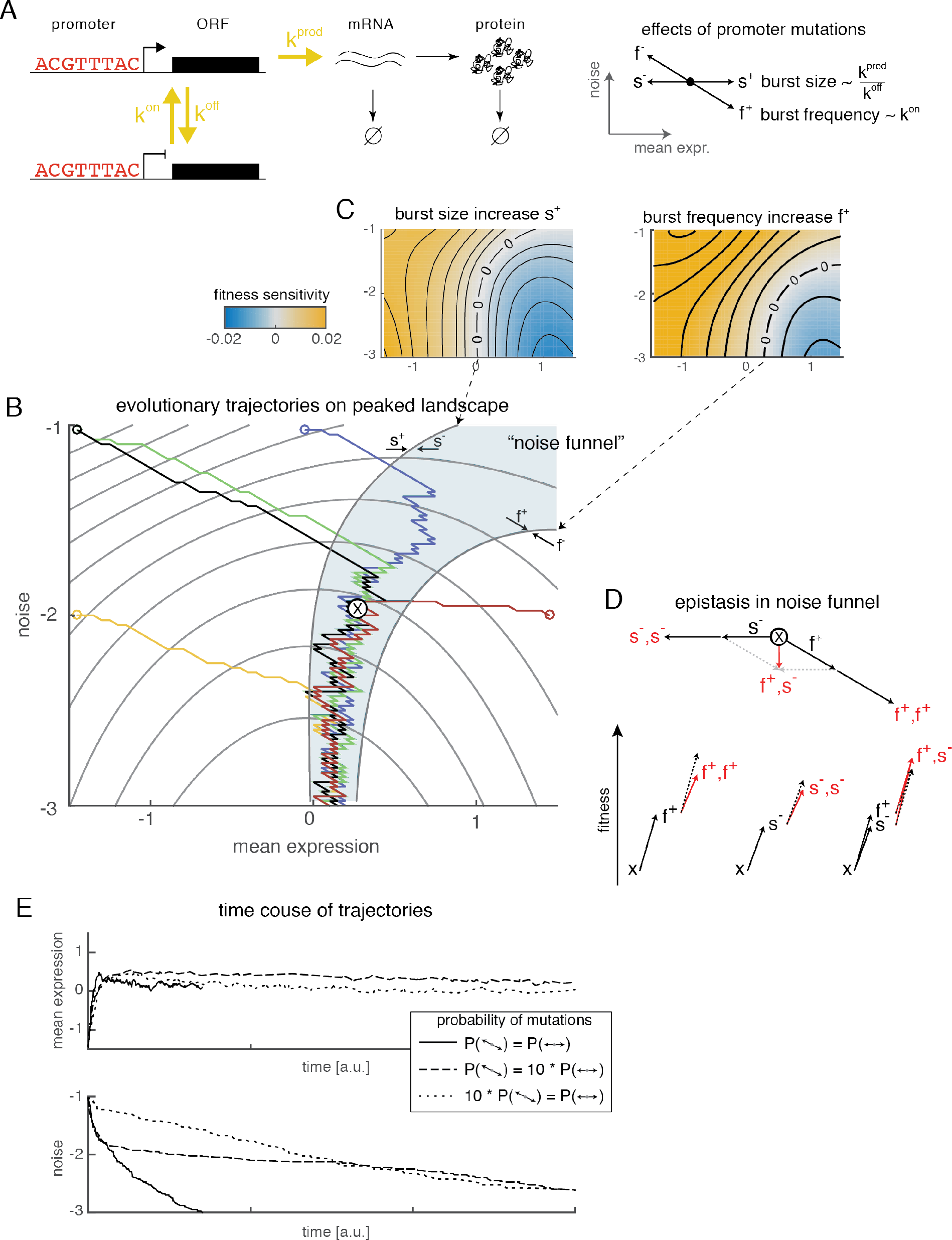
Evolution of optimal gene expression on peaked landscapes proceeds via a noise funnel. (A) Expression of proteins in a two-state telegraph model of transcriptional dynamics. The promoter switches between transcriptionally permissive and prohibitive states, with rates k^on^ and k^off^. In the permissive state mRNAs are transcribed at rate k^prod^, resulting in transcriptional bursts. mRNAs are then translated to proteins; both mRNAs and proteins are degraded eventually. Promoter mutations affect transcriptional bursts by changing their size (by altering k^prod^ or k^off^) or their frequency (by altering k^on^). Mutations that affect burst size result in concordant changes in mean expression, without affecting noise. Mutations that affect burst frequency result in concordant changes in mean expression and opposing changes in noise. (B) Evolution of gene expression on the peaked fitness landscape (sensitivity to protein shortage and surplus) via promoter mutations. Each trajectory is one realization of a stochastic walk, where the likelihood of steps depends on their fitness gains (circles indicate starting genotype). All grid points on the landscape are assumed to be accessible genotype. Shaded area indicates noise funnel, where burst size-decreasing mutations (s^−^) and burst frequency-increasing mutations (f^+^) are beneficial, which results in ratchet-like evolution towards lower noise levels. ‘x’ in noise funnel marks point used to exemplify epistasis in panel (D). (C) Fitness gains of mutations increasing burst size (s^+^, left) or increasing burst frequency (f^+^, right). Borders of no gain (contour marked by ‘0’) do not align due to additional fitness gains from reduced noise in mutations increasing burst frequency, creating the *noise funnel* on the fitness landscape. (D) Upper: Resulting mean/noise effects of consecutive or opposing combinations of increased burst frequency (f^+^) and decreased burst size (s^−^) mutations. Lower: Fitness and epistasis from mutation combinations. First step: fitness of individual mutations. Second step: Expected fitness (black, dashed) and observed fitness (red, solid) of combined mutations. Epistasis is the difference between observed and expected fitness. Starting genotype ‘x’ is marked on landscape in panel (B). (E) Temporal evolution of mean expression and noise on peaked landscape for example trajectories starting at high noise and low fitness (upper left corner in panel (B)). Solid line: Equal probability for burst size and burst frequency mutations to occur, as shown in panel (B). Dashed and dotted lines: Burst frequency or burst size mutations ten times more likely to occur, respectively.

To explore whether the transcriptional process restricts evolutionary trajectories in mean-noise space we simulated stochastic walks on the expression landscapes in which only steps in the directions brought about by promoter mutations are permitted. Here, the next step is probabilistically chosen based on the sensitivities of fitness towards any of the four possible steps (increases or decreases in burst frequency or burst size; step size is always 0.05 log units in mean direction, 0.025 log units in noise direction if changing burst frequency). For simplicity, we assume that mutations affecting burst frequency are as likely to occur as mutations affecting burst size (see Discussion for outcomes of alternative scenarios).

On both principal topologies, we find that the coupling of noise and mean by the transcriptional process restricts the evolution of noise levels. On principal topology 1 (sensitivity to protein shortage) genes evolve towards higher mean expression levels and lower noise levels. The final noise minimum, however, strongly depends on the noise level of the starting point, as noise cannot be further reduced than what is maximally achieved by always selection for frequency increasing over size increasing mutations (Figure 7 - figure supplement 1A). On principal topology 2 (sensitivity to protein surplus) genes evolve towards lower mean expression. Expression noise, however, does at best stay constant (if size altering mutations are selected for) or increases (if frequency altering mutations are selected for), thus moving away from optimally low gene expression noise (Figure 7 - figure supplement 1B). This suggests that, when genes evolve on monotonic, saturating fitness landscapes, the evolution of gene expression noise is fundamentally limited by its coupling to mean expression changes.

In contrast to the principal topologies, evolutionary trajectories on peaked landscapes exhibit a bi-phasic behaviour (Figure 7B). These trajectories are characterized by a first phase of evolution towards optimal mean expression (potentially with coupled changes in expression noise) and a second phase of evolution towards lower expression noise, during which mean expression levels hardly change. Strikingly, independent of the starting point of the simulations, this second phase occurs in a well-defined, narrow region of the landscape (Figure 7B).

We find that this region, which we term the “noise funnel”, is created by a misalignment of the regions where burst frequency and burst size altering mutations are beneficial or detrimental (determined by the points at which equi-fitness lines are tangential to the mutational vectors, Figure 7C). Specifically, here, mutations that *increase* burst frequency and mutations that *decrease* burst size are beneficial, the combination of which results in lowered expression noise but unaltered mean expression (Figure 7D). Consistently, evolution towards lower expression noise in the noise funnel proceeds via alternating steps of increased burst frequency and decreased burst size mutations.

Moreover, evolution towards lower expression noise is accelerated by the epistatic interactions − the non-independence of fitness outcomes - between the two opposing mutations. In particular, a mutation of one type renders a consecutive mutation of the same type less beneficial (Figure 7D), i.e. consecutive mutations of the same type are negative epistatic due to the saturating relationship between fitness and both mean expression and noise (see Figures 4F,G and 6B). The first mutation does, however, render the opposing mutation more beneficial, i.e. their combination is positive epistatic (Figure 7D). The noise funnel therefore acts like an “epistatic ratchet” that uncouples the evolution of noise from mean expression and accelerates the independent minimization of expression noise levels via the genetic interactions of burst size and frequency modulating mutations.

## Discussion

Here we have reconstructed empirical expression-fitness landscapes that allowed us to systematically investigate the quantitative effects of two molecular phenotypes, mean expression and noise, on organismal fitness in yeast.

Across 33 reconstructed landscapes nearly all variance of fitness levels is described by linear combinations of only two principal topologies, which represent the fitness effects of having to too few or too many proteins. These two principal topologies imply that there exist fundamental functional relationships between protein shortage or surplus and organismal fitness that apply to most genes; and that genes only differ in the magnitude of these relationships.

It has been a long-held assumption that genes that are sensitive to sustained depletion or over-expression of their protein abundances are also sensitive to short-lived, stochastic fluctuations in protein abundances (Bar-Even et al., 2006; Batada and Hurst, 2007; Fraser et al., 2004; Keren et al., 2016; Lehner, 2008; Newman et al., 2006). Dedicated experimental tests of this hypothesis, however, had so far remained rare (Duveau et al., 2018), because of the difficulty of independently varying mean expression and noise to quantify the effects of perturbing only one of the two.

Our analyses of how fitness varies across continuous mean-noise fitness landscapes overcomes this limitation, allowing the effects of changes in noise or mean to be examined in isolation as well as in context of each other. This revealed that the more sensitive yeast genes are to changes in their mean abundances the more intolerant are they of high expression noise. Noise intolerance derived from the landscapes predicted endogenous protein expression noise better than did expression-sensitivity, thus suggesting that high expression noise is specifically selected against where it is detrimental to organismal fitness. Moreover, expression fitness landscapes revealed that noise intolerance arises from sensitivities to too few as well as too many proteins and that the fitness impact of non-optimal noise levels is as severe as that of non-optimal mean expression levels.

Our study therefore contributes quantitative empirical evidence for the theory that not only average abundance levels but also short-lived stochastic fluctuations are subject to natural selection in many genes due to their impact on organismal fitness.

Our analysis of expression-fitness landscapes was focused on an eight-fold range around wild-type expression levels, which allowed us to reveal systematic fitness effects across many genes. The fitness effects of expression noise are, however, expected to depend on the discrepancy between the actual and optimal average expression levels (Tanase-Nicola and ten Wolde, 2008). Especially, high expression noise should become beneficial when average expression is far away from optimum, as this would allow few cells to transiently express more optimal protein abundances, therefore increasing overall population growth rate, and this has recently been demonstrated for the TDH3 gene in yeast (Duveau et al., 2018). Indeed, when examining expression fitness landscapes initially excluded from our analyses (due to wild-type expression levels outside of the investigated mean expression range), we find examples of this transition in noise-fitness effects. For the two highly expressed genes ENO2 and RPL3, high expression noise turns from being detrimental when mean expression is close to wild-type levels to beneficial when mean expression drops far below wild-type expression levels (Figure 7 - figure supplement 2). The fitness at low mean expression and high noise is, however, much lower than fitness at more optimal mean expression and low noise. This shows that, while high noise can improve fitness if expression is far from its optimum, it is by no means a substitute for optimally tuned expression levels (Wolf et al., 2015).

We have further used the concept of expression fitness landscapes to study evolutionary scenarios for gene expression. The expression fitness landscapes revealed that low noise levels should accelerate the evolution of optimal mean expression levels. Therefore, when expression is close to its optimum, low noise not only maximizes fitness but can also enhance evolvability (Lehner, 2008; Peterson et al., 2009).

In contrast, selection pressure on expression noise is strongest when mean expression levels are optimal, especially when genes are sensitive to both protein shortage and surplus (‘peaked’ landscapes). Notably, this is consistent with findings for the TDH3 gene in yeast - which has a ‘peaked’ fitness profile (Duveau et al., 2017) -, where noise-increasing but not mean-altering mutations are under purifying selection (Metzger et al., 2015).

However, how can noise levels evolve if mean expression is already optimal, given that due to protein production constraints noise can only change in concert with mean expression? Investigation of evolutionary scenarios explicitly accounting for these phenotypic constraints revealed that combined sensitivities to protein shortage and surplus create a narrow landscape region − the noise funnel − in which the evolution of noise is uncoupled from mean expression. The noise funnel is the consequence of a disagreement in the signs of fitness effects of burst size and burst frequency modulating mutations. The independent evolution of low noise levels is further promoted by an epistatic ratchet mechanisms in which the combination of both mutation types are positive epistatic but two consecutive mutations of the same type are negatively epistatic.

We performed these evolutionary simulations under the simplifying assumption that burst size and burst frequency mutations are equally likely to occur. Typically, mutations that change burst frequency are, however, much more likely to occur in promoter regions than mutations affecting burst size (Hornung et al., 2012). Consistent with the ratchet-like epistatic interplay of both mutations types in the noise funnel, we find that an equal likelihood for both types of mutations to occur is key to rapid reduction of expression noise within the noise funnel (Figure 7E). Evolution of minimal gene expression noise would therefore be hampered if burst size could only change via mutations in the promoter. Changes in post-transcriptional processes, however, also affect the size of expression bursts (Pedraza and Paulsson, 2008), thus enlarging the mutational target space for burst size changing mutations and potentially accelerating the evolutionary minimization of expression noise level. The vast expansion of post-transcriptional repressive regulators in higher eukaryotes, such as microRNAs, could therefore facilitate the reduction of gene expression noise across distinct cellular states (Bartel and Chen, 2004; Peterson et al., 2009; Schmiedel et al., 2015). Consistently, human dosage-sensitive genes are highly enriched for microRNA binding sites (Schmiedel et al., 2017).

Together, this shows that in order to understand the evolution of gene expression both the constraints imposed by the underlying molecular mechanisms as well as the mapping between expression distributions and organismal fitness have to be considered. Moreover, our analysis makes the testable prediction that for ‘peaked’ genes, regulatory elements with opposing influences on burst size and burst frequencies should co-evolve in order to minimize expression noise.

## Methods

### Data acquisition

| Data | Source |
| --- | --- |
| Mean expression levels of endogenous gene promoter and synthetic promoters when driving YFP | (Keren et al., 2016) |
| Read counts of gene-promoter combinations in competitive growth experiments on glucose (used to calculate strain fitness) | Gene Expression Omnibus accession number GSE83936 (Keren et al., 2016) |
| Mean expression and noise levels of synthetic promoters driving YFP | (Sharon et al., 2014) |
| Synthetic promoter sequences, list of transcription factor motives | (Sharon et al., 2012) |
| Protein expression noise of GFP-fusions in haploid cells | (Newman et al., 2006) |
| Protein expression noise of GFP-fusions in diploid cells | (Stewart-Ornstein et al., 2012) |
| Haplo-insufficient yeast genes | (Deutschbauer et al., 2005) |
| Over-expression sensitive yeast genes | (Makanae et al., 2013; Sopko et al., 2006) |
| Essential yeast genes | Saccharomyces Genome Deletion Project (Giaever et al., 2002)<br>http://www-sequence.stanford.edu/group/yeast_deletion_project/Essential_ORFs.txt |

### Data processing

#### Fitness

Relative fitness for growth in glucose of each promoter-gene pair strain was calculated from changes in read count frequencies across the competitive growth experiment (Keren et al., 2016). Fitness at two time-points (23h and 35h growth) were calculated as 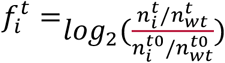, with *n* as the number of reads (supplemented with a pseudo count of 0.1), subscripts denoting strain *i* or the wildtype *wt*, and superscripts denoting the time-point (*t0* as starting time-point of the competition experiment, *t* for the two later time-points). A linear model was fit to derive a normalization factor to correct systematic fitness differences across all promoter-gene pair strains between the two time-points. Fitness for each promoter-gene pair was then calculated as the weighted average of relative fitness measures at both time-points, with weights as the inverse of error estimates calculated from read counts as 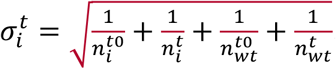. A combined error of fitness for each promoter-gene pair was accordingly derived as 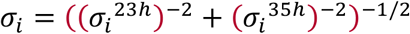.

#### Promoter expression properties

Promoter mean expression and promoter expression noise was calculated as average over two replicates (Sharon et al., 2014). Error of both measures was estimated as the running average of replicate standard deviation as a function of sequencing read based error estimate over all promoters (calculated using MATLAB function *fit*, with method *loess* and span 0.5)

#### Quality control

Promoters were checked for consistency of mean expression estimates between driving YFP on a plasmid (Sharon et al., 2014) and driving YFP from the HIS3 locus (Keren et al., 2016). A linear model fit to the log_2_-transformed mean expression data was used to transform the plasmid-derived data in order to make the two studies comparable. 11 of 120 promoters that showed a log_2_-derivation of more than 0.5 between mean expression estimates in both studies were discarded (Figure 2 - figure supplement 1A). Another 6 promoters that had a median fitness error estimate over all promoter-gene combinations greater than 0.1 were discarded (Figure 2 - figure supplement 1B). Finally, to restrict our analysis to a sufficiently homogenously populated core region in the mean-noise space, 24 promoters with mean expression below 2 or above 6 log_2_-expression units were discarded (Figure 2 - figure supplement 1C). Because our subsequent analyses are focused on the fitness effects around the wild-type expression of genes, only those 33 of 85 genes that have an estimated mean expression output of their wild-type promoters that lies in the centre of the analysed expression range (between 3 and 5 log_2_-expression units) were considered (Figure 2 - figure supplement 1D). Additionally, for the transcription factors ABF1, MIG1 and RAP1 several promoter-gene pairs (3, 5 and 2, respectively) were discarded from our analysis because the promoters contain predicted binding motifs for these genes.

### Calculation of noise-mean fitness landscapes

To reconstruct a smooth, continuous fitness landscape for each gene, we calculated fitness values on a regular grid across the mean-noise space using a Gaussian smoothing approach. The grid dimensions were chosen such that the rectangular grid covers all promoter strains in the mean noise space and grid points were spaced by 0.05 log_2_-mean expression units and 0.025 log_2_-noise (CV^2^) units. For subsequent analyses of expression fitness landscape features, we investigated grids that extend ±1.5 log_2_-mean expression units from the wild-type promoter expression and range between −3 and −1 log_2_-noise units (see below) and thus have 61×81=4941 grid points. For visualization purposes (Figure 2F) we also computed more extensive grids. For each gird point *xy*, a fitness value was calculated as the weighted average over the fitness of all gene-specific strains. How the strain in which promoter *i* drives the gene *j* contributes to fitness at grid point *xy* was calculated by integrating over the joint probability density function of a Gaussian smoothing kernel centred on the grid point and a Gaussian likelihood function centred on the promoter position in mean-noise space. The Gaussian smoothing kernel is a bi-variate normal density (Matlab function *mvnpdf*) with means *μ* = *x* and η = *y*, the grid point position in mean-noise space, and covariance matrix 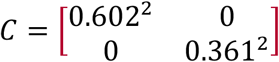, the optimal shape of the kernel that minimizes the RMSE between fitness surfaces and measured fitness of promoter-gene strains, as estimated from ten-fold cross validation. The Gaussian likelihood function of the true position of the promoter *i* in mean-noise space is a bi-variate normal density with means *μ* = *μ_i_* and *η* = *η_i_*, the estimates of mean expression and noise of the promoter, and covariance matrix 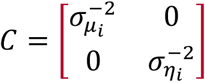, the error estimates for mean expression and noise of the promoter. The integral over the joint probability densities, further normalized by the uncertainty of the fitness estimate of promoter-gene strain *ij*, results in the weighting of the fitness of promoter-gene strain *f_ij_* for the fitness at grid point *xy* in the fitness landscape of gene *j*.

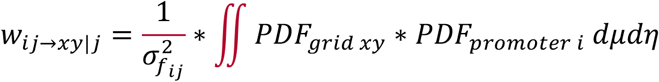

In practice, to speed up computations at little cost to precision, *w*_*ij*→*xy*|*j*_ was calculated on a 21×21 auxiliary grid around the grid point *xy*, with spacing 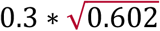 in the mean and 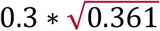 in the noise expression direction and only using those auxiliary grid points where the smoothing kernel probability density is larger than 1% of the respective density on the grid point *xy*.

The fitness at grid point *xy* in the fitness landscape of gene *j*, *f*_*xy*|*j*_, is the weighted average over the fitness values of all gene-specific strains:

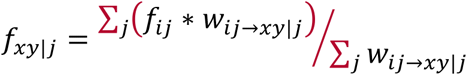

Note that some landscapes have non-optimal fitness at wild-type expression, especially steeper landscapes with asymmetric shapes (Figure 3), such as RPN8 (Figure 2F). These effects might result from two plausible causes that are both due to blurring of fitness landscapes: first, the noise levels assayed by the synthetic promoters might be higher than the noise levels of the endogenous promoters, i.e. at even lower noise levels the actual expression-fitness relationships are sharper and fitness at wild-type expression is optimal, or second, the resolution limit of our smoothing procedure, which in turn is dictated by the experimental errors in the data. We conclude that the observed effect should not impact the generality of our results.

### Principal component analysis of fitness landscapes

To understand whether the expression fitness landscapes share common topological features we performed a principal component analysis (PCA) across all landscapes (Figure 4 - figure supplement 1). For this analysis, landscapes extending ±1.5 log_2_-mean expression units from each gene’s wild-type promoter expression and ranging from -3 to-1 log_2_-noise units were compared between the 33 genes. Prior to performing the PCA, fitness values on each landscape were normalized to the fitness at wild-type expression and log_2_-noise = −3. PCA was performed with Matlab function *pca* (option *‘Centered’* set to *true*) using the fitness on each of the 61×81=4941 grid points of each gene’s landscape as observations and treating the 33 genes as variables. Reported principal component loadings (‘principal topology loadings’) for each landscape were corrected for the loadings of the mean fitness landscape (its loadings for the first and second components are 1.03 and 0.15, respectively).

### Expression sensitivity and noise intolerance

#### Calculation and comparison of metrics

Expression sensitivity of each gene was calculated as the average curvature (second derivative) of the mean expression-fitness function at log2-noise = -3. Noise intolerance of each gene was calculated as the maximal negative slope (first derivative) of the noise-fitness function at wild-type mean expression. Expression sensitivity and noise intolerance were also computed for 1.000 randomizations of each gene’s fitness landscape, where in each randomization the fitness values between all promoter-gene strains were permutated. P-values for each gene’s expression-sensitivity and noise intolerance were calculated as the fraction of the gene’s randomized fitness landscapes with expression-sensitivity and noise intolerance values greater or equal to the non-randomized values. Positive false discovery rate was calculated from these p-values using the linear step-up procedure (Matlab function *mafdr* with option ‘BHFDR’). Additionally, the Pearson correlation between expression sensitivity and noise intolerance of all genes was calculated for each randomization run. A p-value for the Pearson correlation coefficient between expression sensitivity and noise intolerance on real landscapes was derived as the fraction of correlation coefficients from randomization runs that are greater or equal than that of the real data.

Expression-sensitivity and noise intolerance were also derived from ‘raw’ gene-promoter strain data. Here, for each gene the Pearson partial correlation coefficients between noise or mean expression levels and fitness of gene-promoter strains were calculated while controlling for the other expression phenotype (using Matlab function *partialcorr*). P-values for alternative hypothesis that partial correlation is not 0 were used to calculate positive false discovery rate using the linear step-up procedure (Matlab function *mafdr* with option ‘BHFDR’). As for the rest of our analysis we only considered promoters within the expression range of 2 to 6 log_2_-mean expression units (Figure 2 - figure supplement 1C). Not unexpectedly, correlation between expression-sensitivity and noise intolerance are somewhat smaller, which might stem from the fact that partial correlations can only identify linear dependencies, but e.g. not the peaked expression-fitness relationships of many landscapes.

Moreover, we used two additional ‘expression-sensitivity’ measures from published data. First, the ‘expression curvature’ metric used by Keren et al. (2016) was calculated as described therein as the minimal mean expression distance at which a 5% fitness drop compared to fitness at wild-type expression is observed (on impulse fitted fitness data as reported in Supplementary Table S3 of (Keren et al., 2016)). Second, we derived a ‘dosage sensitivity’ metric, where genes were considered dosage sensitive if they have been reported to be essential (Giaever et al., 2002) (n = 23; n = 3 of which are also haplo-insufficient (Deutschbauer et al., 2005)) or over-expression sensitive (Makanae et al., 2013; Sopko et al., 2006) (n = 11, nine of which are also essential) in large-scale genetic screens.

#### Comparison to endogenous noise levels

Noise intolerance and the three metrics of expression sensitivity of genes were compared to endogenous noise levels reported in large-scale screens by calculating the Spearman rank correlation coefficient. P-values were derived for the alternative hypothesis that correlation is smaller than 0 and aggregated for the three tests of each metric using Fisher’s method. Endogenous noise levels in haploid cells when grown in minimal medium (SD) or rich medium (YPD) for 18 and 22 genes out of the 33 genes investigated here were obtained from Newman et al. (2006). Reported noise DM values (deviation from running median) were used for comparison. Additionally, endogenous noise levels in diploid cells for 9 out of the 33 genes were obtained from Stewart-Ornstein et al. (2012). Noise levels in diploid were similarly corrected for the running median of noise levels across expression levels.

### Fitness sensitivities across landscapes

Fitness sensitivities to changes in noise or mean expression were calculated as the gradient between adjacent grid points in the respective directions (using Matlab function *gradient*) and normalized to represent fitness changes upon a two-fold change in the expression phenotypes. Equi-fitness lines were determined with Matlab function *contourc* at relative fitness values in the range of [-0.05,0.025] with a spacing of 0.005. Only fitness sensitivities on points across these equi-fitness lines are shown in Figure 6.

### Evolution on fitness landscapes

Evolutionary simulations on fitness landscapes in a promoter mutation scenario were implemented as stochastic walks using a Gillespie algorithm (Gillespie, 1977). Here, the probability of a mutation to be selected for is proportional to its fitness gain relative to the summed fitness gains from all mutations with non-negative fitness gains. The time until the next mutation is selected for exponentially distributed with mean proportional to the inverse sum of non-negative fitness gains. Each step results in the jump to an adjacent grid point. For burst size mutations, this jump is to an adjacent grid point with altered mean expression (plus or minus 0.05 log_2_-mean expression units, if an increase or a decreased burst size is selected for, respectively) but equal noise. For burst frequency mutations to grid points with both altered mean expression (plus or minus 0.05 log_2_-mean expression units, if an increase or a decreased burst frequency is selected for, respectively) and altered noise (minus or plus 0.025 log_2_-mean expression units, respectively; grid is spaced twice as narrow in noise direction, thus the change in noise for a burst frequency mutation is the negative square root change in mean expression).

To simulate differential likelihoods of mutations (related to Figure 7E), we modified the Gillespie algorithm by altering the calculation of probabilities for mutational selection and time intervals. E.g. for the scenario where burst size mutations are ten times less likely, their fitness gains were divided by a factor of ten in the calculation of probabilities, i.e. they were ten times less likely to be selected for.

## Code availability

All analysis was performed using Matlab version R2014b. All code and source data can be found at https://github.com/jschmiedel/NMF_landscapes.

## Acknowledgements

This work was supported by a European Research Council Consolidator grant (616434), the Spanish Ministry of Economy and Competitiveness (BFU2011-26206 and SEV-2012-0208), the AXA Research Fund, Agència de Gestió d’Ajuts Universitaris i de Recerca (AGAUR, 2014SGR831), FP7 project 4DCellFate (277899), the EMBL-CRG Systems Biology Program (all to B.L.), an EMBO Long-Term Fellowship (ALTF 857-2016), the European Union’s Horizon 2020 research and innovation programme (Marie Skłodowska-Curie grant agreement No 752809) (both to J.M.S) an AGAUR grant (2014SGR0974) and a MINECO grant (BFU2015-68351-P) (both to L.B.C.).

## Author Contributions

JMS performed all analyses. JMS, LBC and BL conceived the study. JMS and BL designed analyses and wrote the manuscript with input from LBC.

---

**Figure 2 - figure supplement 1.**
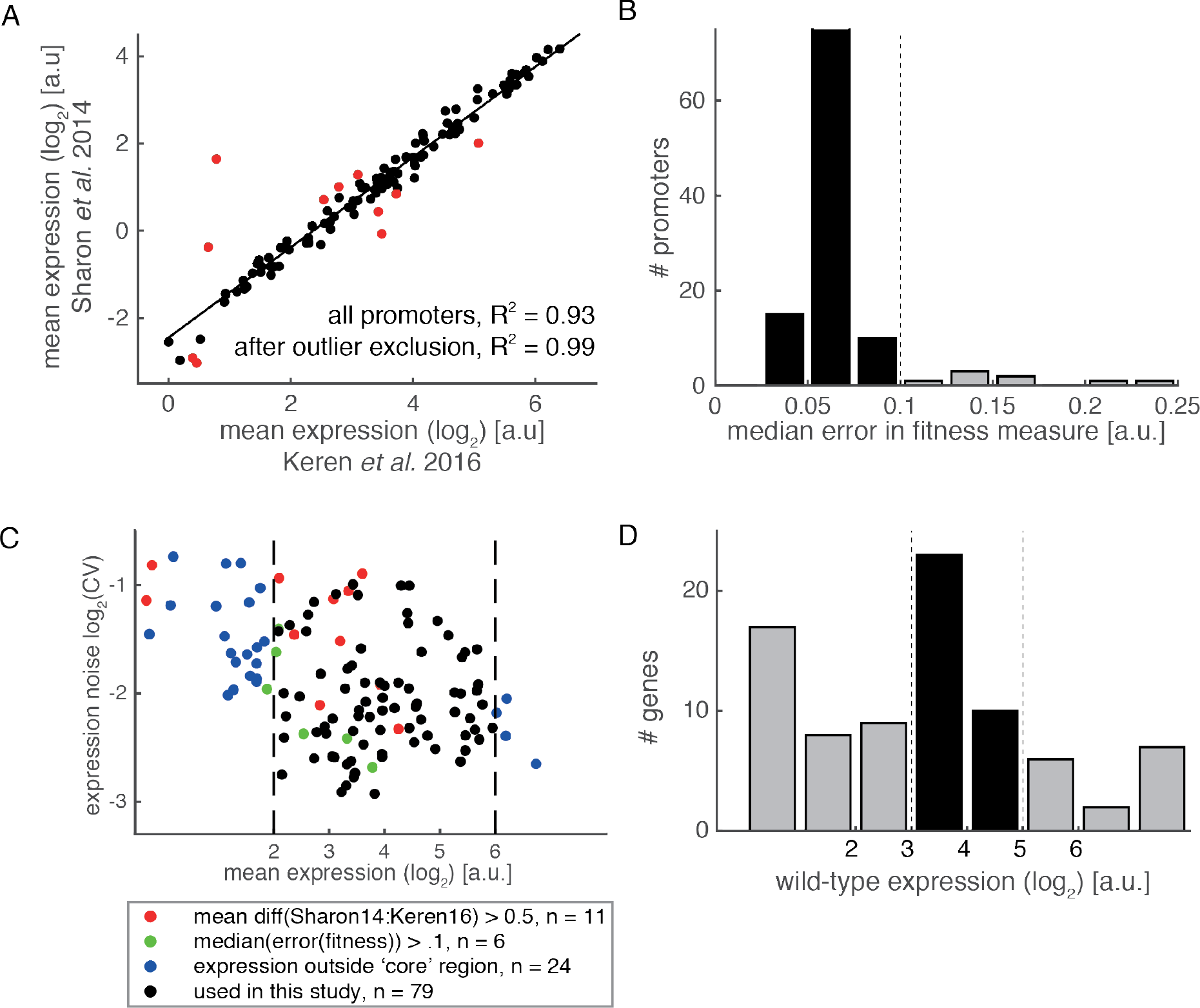
Processing of experimental data. (A) Relative mean expression strength of synthetic promoters when driving a genomically integrated YFP measured for individual strains (Keren et al., 2016) versus when driving a plasmid-based YFP measured in a pooled fashion with a fluorescent activated cell sorting and sequencing approach (Sharon et al., 2014). Outliers excluded from further analysis (because their deviation is greater than 0.5 log2-units from proportionality relationship) are marked in red. Squared Pearson correlation coefficients and corresponding p-values for correlation coefficients being zero are indicated. (B) Median error of fitness estimates for each promoter across all 85 genes. Promoters with median errors greater than 0.1 were discarded from analysis. (C) Promoters in mean-noise expression space. Promoters excluded because of too large discrepancies in mean expression strengths between the two studies (panel A) are marked in red (n = 11). Promoters with too high errors of fitness measurements (panel B) are marked in green (n = 6). Promoters that lie outside of the core mean-noise space region analysed (mean range [2,6] log2-units) are marked in blue (n = 24). The remaining 79 promoters used in this study (black) are homogenously distributed in the analysed mean-noise space. (D) Estimated wild-type expression strength of endogenous promoters of the 85 investigated in Keren et al. (2016). We focused on genes whose promoters have an expression strength in the centre (3 to 5 log2 expression units) of the mean-noise space region analysed.

**Figure 4 - figure supplement 1.**
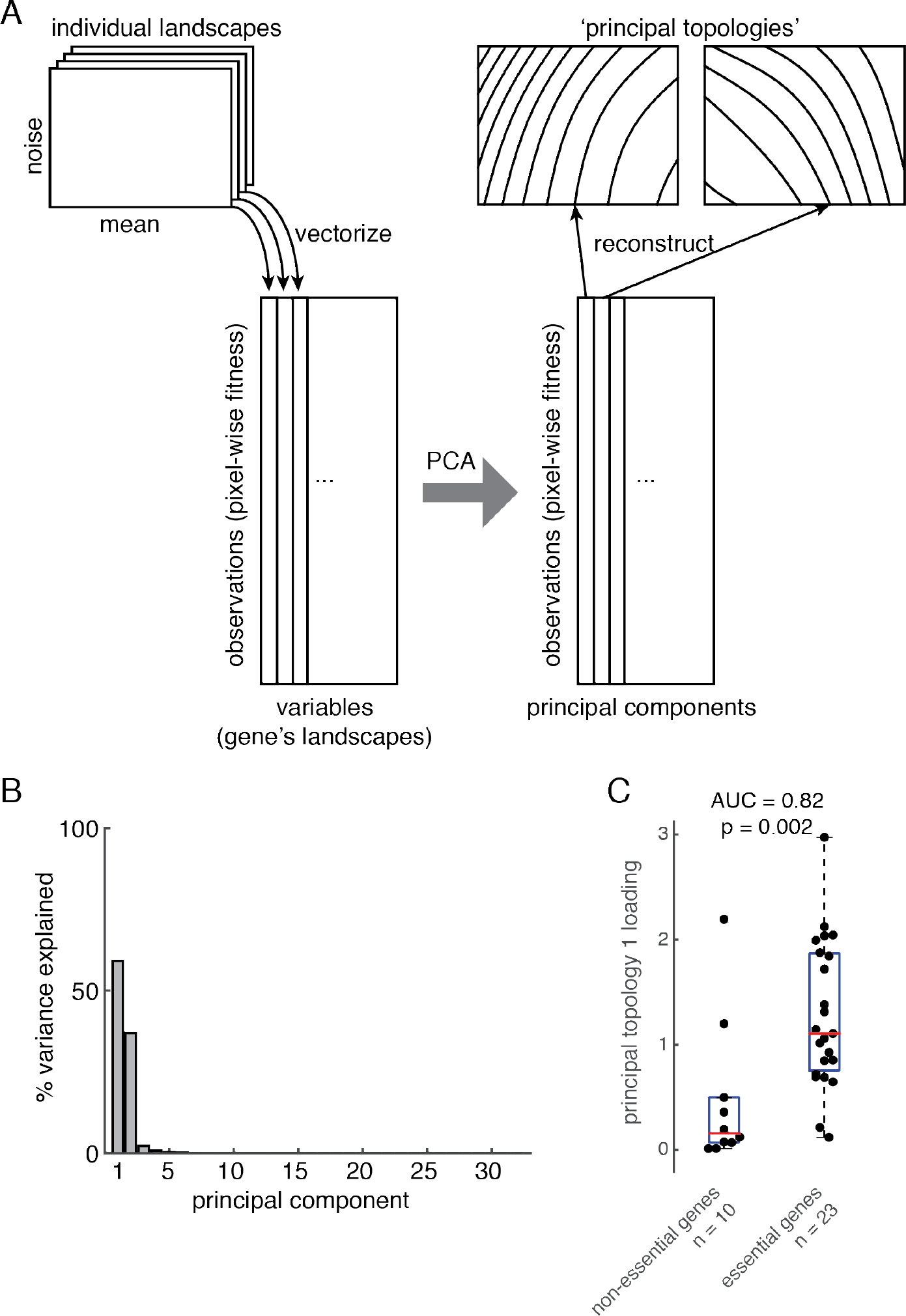
Principal component analysis of expression-fitness landscapes. (A) Overview of principal component analysis. Individual landscapes are vectorised and concatenated into an N x L matrix, with N the number of grid points on the landscapes and L the number of landscapes (33). Principal component analysis on this matrix yields a matrix of principal components, the first two (explained the most variance on individual landscapes) from which ‘principal topologies’ are reconstructed. (B) Percent variance of fitness landscapes explained by the principal components. (C) Principal topology 1 loadings of genes correspond with genes’ essentialities as assessed from gene deletion screens. Area under the curve (AUC) of receiver operating characteristic (Matlab function *perfcurve*) as well as one-sided p-value from Wilcoxon rank sum test (Matlab function *ranksum*) are indicated.

**Figure 5 - figure supplement 1.**
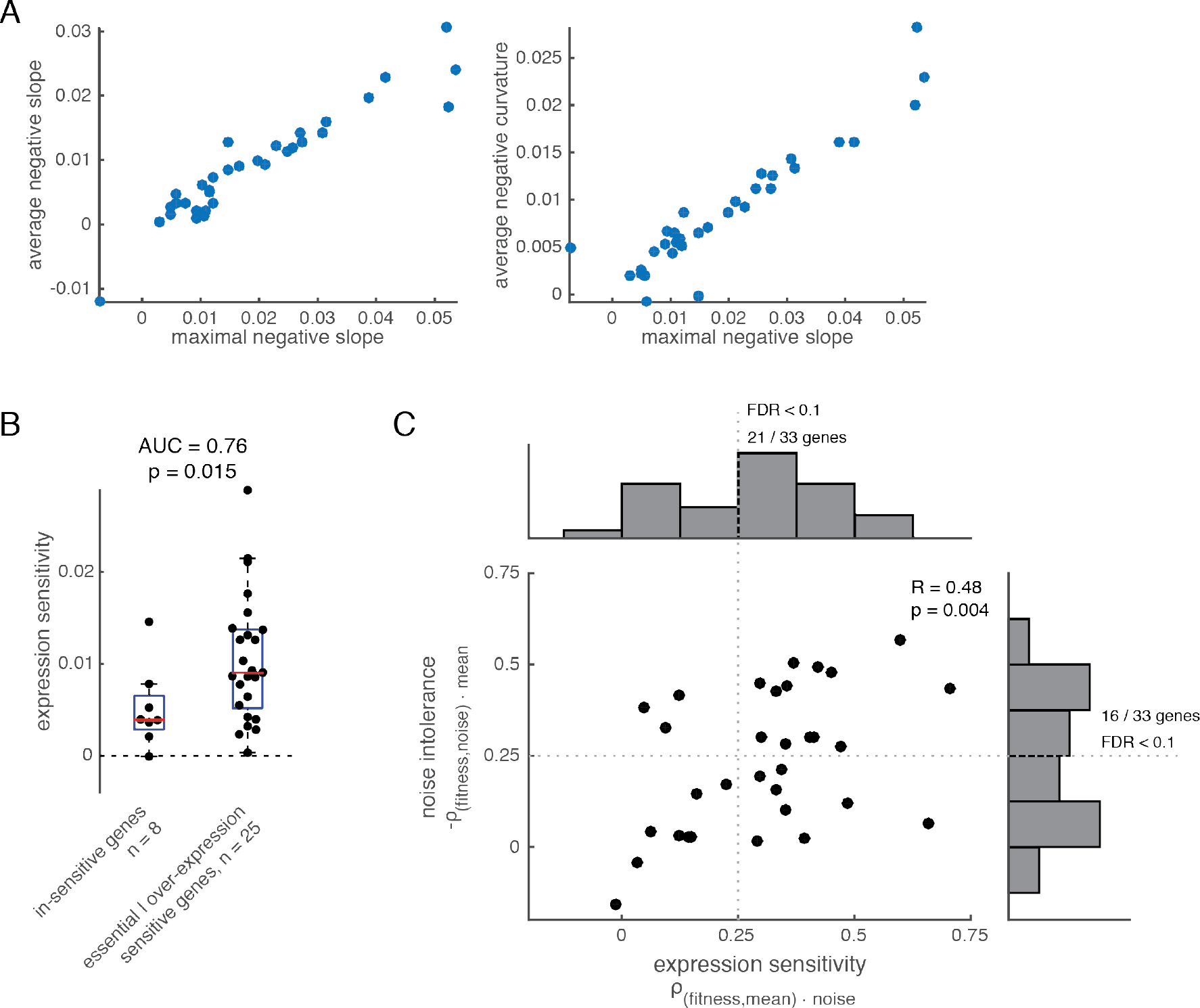
Expression sensitivity and noise intolerance on expression-fitness landscapes. (A) Comparison of metrics to calculate noise intolerance on individual expression-fitness landscapes. ‘Maximal negative slope’ is the metric described and used in the main text (maximal negative slope of noise-fitness function at wild-type mean expression). Similarly, ‘average negative slope’ is the average of the negative slope of the noise-fitness function at wild-type mean expression. ‘Average negative curvature’ is the average of the negative second derivate of the noise-fitness function at wild-type mean expression. (B) Expression sensitivity of known dosage sensitive (essential and/or over-expression sensitive) genes. Area under the curve (AUC) of receiver operating characteristic (Matlab function *perfcurve*) as well as one-sided p-value from Wilcoxon rank sum test (Matlab function *ranksum*) are indicated. (C) Relationship between expression sensitivity and noise intolerance across all genes when derived from Pearson partial correlation of mean or noise with fitness values while controlling for the other expression phenotype from raw gene-promoter strain data. False discovery rate (FDR) < 10% threshold indicated in both dimensions (Matlab function *mafdr* using Benjamini-Hochberg correction). Histograms on top and to the right show distributions of genes across expression sensitivity and noise intolerance, respectively. Indicated are the numbers of genes with FDR < 10%. Pearson correlation coefficient of expression sensitivity and noise intolerance across genes and p-value are indicated.

**Figure 7 - figure supplement.**
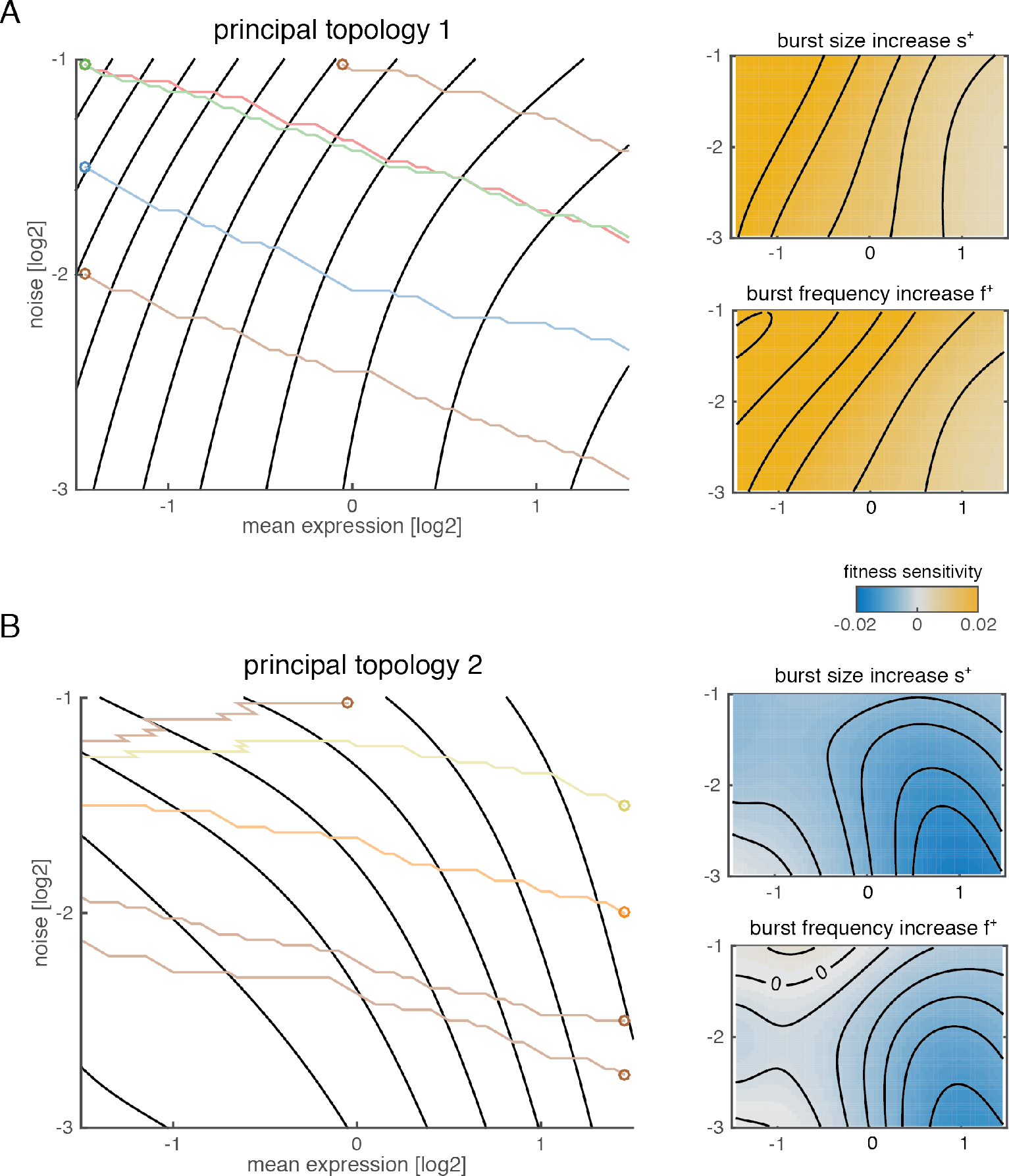
Evolution of gene expression on principal topologies. Left: Evolution of gene expression on principal topology 1 (sensitivity to protein shortage, panel **A**) or principal topology 2 (sensitivity to protein surplus, panel **B**) via promoter mutations. Each trajectory is one realization of a stochastic walk, where likelihood of steps depends on their fitness gains (circle indicates start point). All grid points on the landscape are assumed to be accessible genotype. Right: Fitness gains of mutations increasing either burst size (s^+^, left) or burst frequency (f^+^, right) across the corresponding landscapes.

**Figure 7 - figure supplement 2.**
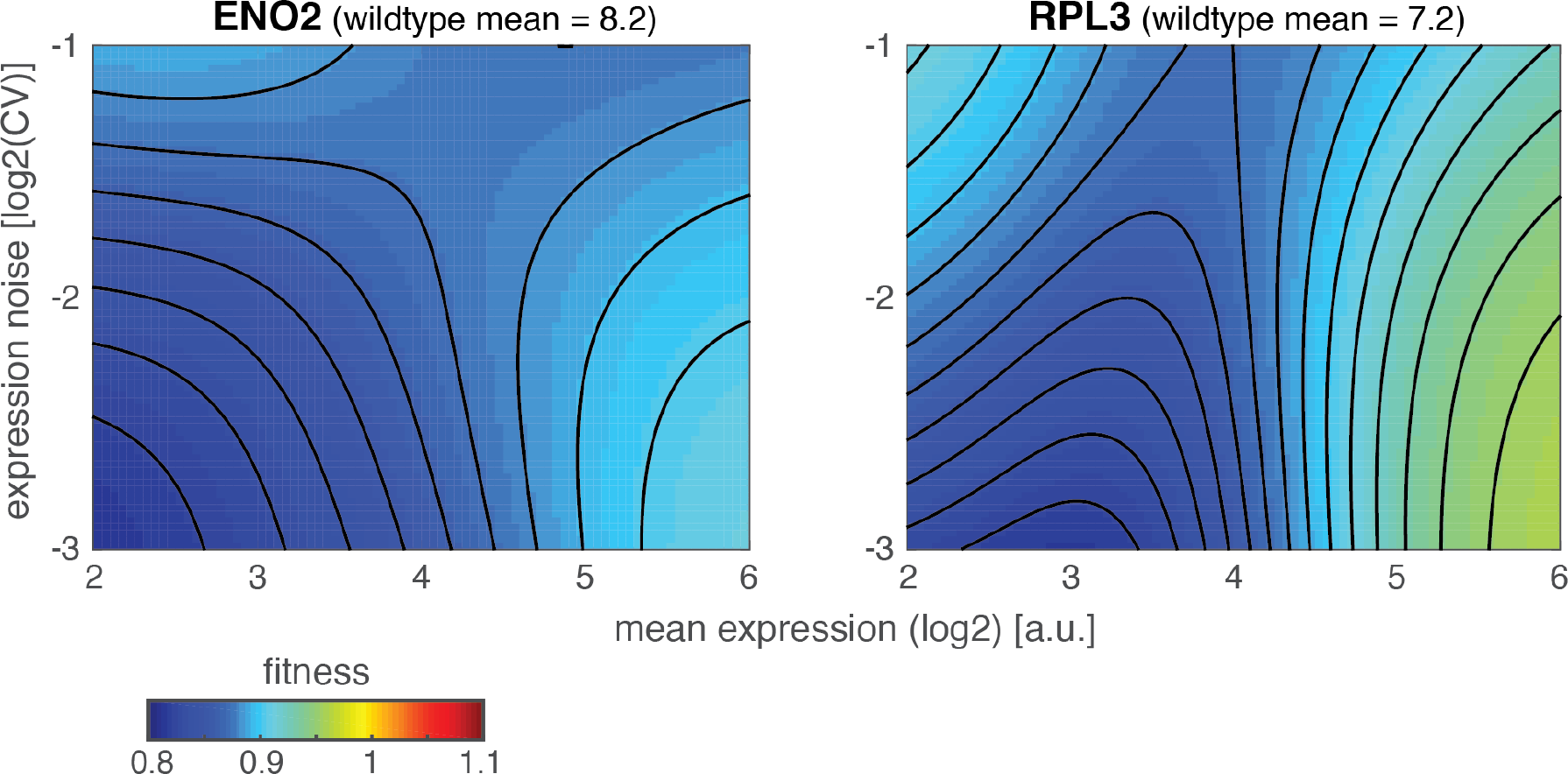
Reversal of noise-fitness effects far away from optimal mean expression. Expression-fitness landscapes of two genes (ENO2, wildtype mean expression = 8.2 log2-units; RPL3 wildtype mean expression = 7.2 log2-units) for which high expression noise transitions from being detrimental close to wild-type mean expression (mean expression above ~5 log2-units) to beneficial far below wild-type mean expression (mean expression below ~3.5 log2-units).

